# Exposing flaws in S-LDSC; reply to Gazal *et al*.

**DOI:** 10.1101/280784

**Authors:** Doug Speed, David J Balding

## Abstract

In our recent publication,^1^ we examined the two heritability models most widely used when estimating SNP heritability: the GCTA Model, which is used by the software GCTA^2^ and upon which LD Score regression (LDSC) is based,^3^ and the LDAK Model, which is used by our software LDAK.^4^ First we demonstrated the importance of choosing an appropriate heritability model, by showing that estimates of SNP heritability can be highly sensitive to which model is assumed. Then we empirically tested the GCTA and LDAK Models on GWAS data for a wide variety of complex traits. We found that the LDAK Model fits real data both significantly and substantially better than the GCTA Model, indicating that LDAK estimates more accurately describe the genetic architecture of complex traits than those from GCTA or LDSC.

Some of our most striking results were our revised estimates of functional enrichments (the heritability enrichments of SNP categories defined by functional annotations). In general, estimates from LDAK were substantially more modest than previous estimates based on the GCTA Model. For example, we estimated that DNase I hypersensitive sites (DHS) were 1.4-fold (SD 0.1) enriched, whereas a study using GCTA had found they were 5.1-fold (SD 0.5) enriched,^5^ and we estimated that conserved SNPs were 1.3-fold (SD 0.3) enriched, whereas a study using S-LDSC (stratified LDSC) had found they were 13.3-fold (SD 1.5) enriched.^6^

In their correspondence, Gazal *et al.* dispute our findings. They assert that the heritability model assumed by LDSC is more realistic than the LDAK Model, and that estimates of enrichment from S-LDSC^7^ are more accurate than those from LDAK. Here, we explain why their justification for preferring the model used by LDSC is incorrect, and provide a simple demonstration that S-LDSC produces unreliable estimates of enrichment.

## The GCTA and LDAK Models

Let 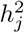 denote the heritability contributed by SNP *j*, defined so that 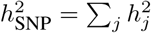 is the SNP heritability of the trait. The GCTA Model assumes a prior distribution for effect sizes such that each SNP is expected to contribute equal heritability: 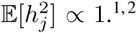.^1,2^ By contrast, the LDAK Model assumes

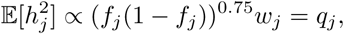

where *f*_*j*_ is the minor allele frequency (MAF) of SNP *j* and *w*_*j*_ is its LDAK weight (SNPs in regions of high LD tend to have lower *w*_*j*_, and vice versa).^1, 4^ If 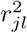 denotes the squared correlation between SNPs *j* and *l*, then 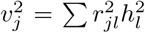 is the total heritability tagged by SNP *j* (in theory, the summation is across all SNPs, but in practice^3^ we consider only those within 1 cM). Under the GCTA Model, 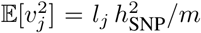, where 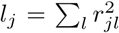 is the LD score of SNP *j*,^3^ whereas under the LDAK Model, 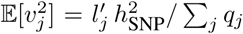, where 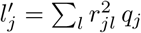 is the “LDAK score” of SNP *j*.

## LDSC is based on the GCTA Model

Suppose we have a GWAS on *n* individuals and *m* SNPs. The *χ*^2^(1) additive association test statistic for SNP *j* has value^8^

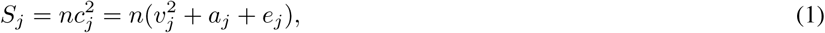

where 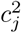 is the phenotypic variance explained by SNP *j*, which can be partitioned into 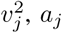 and *e*_*j*_, components corresponding to causal variation, confounding and noise, respectively. *e*_*j*_ has expectation 1*/n*; LDSC seeks to estimate the expected values of 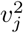 and *a*_*j*_. For this it assumes the model^3^

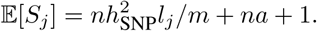

We can see that this follows from Equation (1) if we assume the GCTA Model (as then 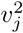 has expected value 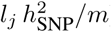) and that *a*_*j*_ is constant across the genome.

## Evidence for the LDAK Model

In our previous work,^1^ we performed a careful evaluation of the GCTA and LDAK Models. We collected GWAS data for 42 different traits, both binary and quantitative, then performed stringent quality control, checking that any confounding due to population structure or cryptic relatedness was at most slight.^9, 10^ We demonstrated that it was valid to compare models using the REML likelihood, then used this approach to show that the LDAK Model was both significantly and substantially more realistic than the GCTA Model; it fit better for 37 of the 42 traits (*P <* 10^−7^) and resulted in an average increase in log likelihood of 9.8 per trait. We also investigated attempts to improve the accuracy of the GCTA Model by partitioning (we focused on GCTA-LDMS,^11^ but the same arguments apply to S-LDSC^7^). While partitioning allowed GCTA to achieve log likelihoods comparable to those from LDAK, this came at the cost of 19 extra parameters which were arbitrarily defined, added little to model interpretation and reduced the precision of heritability estimates.

## Evidence for the GCTA Model

In their correspondence, Gazal *et al.* make no mention of the evidence we provided in support of the LDAK Model. Instead, their rationale for preferring the GCTA Model is the observation that for many traits *the marginal effect size of a SNP has been shown to have a strong linear dependency on its LD score* (in our notation, that there is a significant correlation between *l*_*j*_ and *S*_*j*_). We do not dispute that these correlations exist; for example, Figures 1a & 1b demonstrate that *l*_*j*_ and *S*_*j*_ are correlated for human height, using data from the most recent Giant Consortium meta-analysis.^12^ However, we disagree with the reasoning that because the GCTA Model predicts 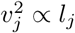, an observed correlation between *l*_*j*_ and *S*_*j*_ is evidence to prefer the GCTA Model. Firstly, it does not immediately follow from Equation (1) that all correlation between *l*_*j*_ and *S*_*j*_ is driven by correlation between *l*_*j*_ and 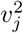. While this would be true if *a*_*j*_ = *a* (or more generally, if *a*_*j*_ is orthogonal to *l*_*j*_), no empirical evidence was provided to support this assumption.^3^ Considering that *l*_*j*_ correlates with factors such as MAF, genotyping certainty and population axes (Supplementary Figure 1), it seems plausible that *a*_*j*_ does correlate with *l*_*j*_. It was to avoid uncertainty regarding *a*_*j*_, that when comparing the GCTA and LDAK Models,^1^ we restricted ourselves to GWAS where we were confident that *a*_*j*_ ≈ 0.

**Figure 1:**
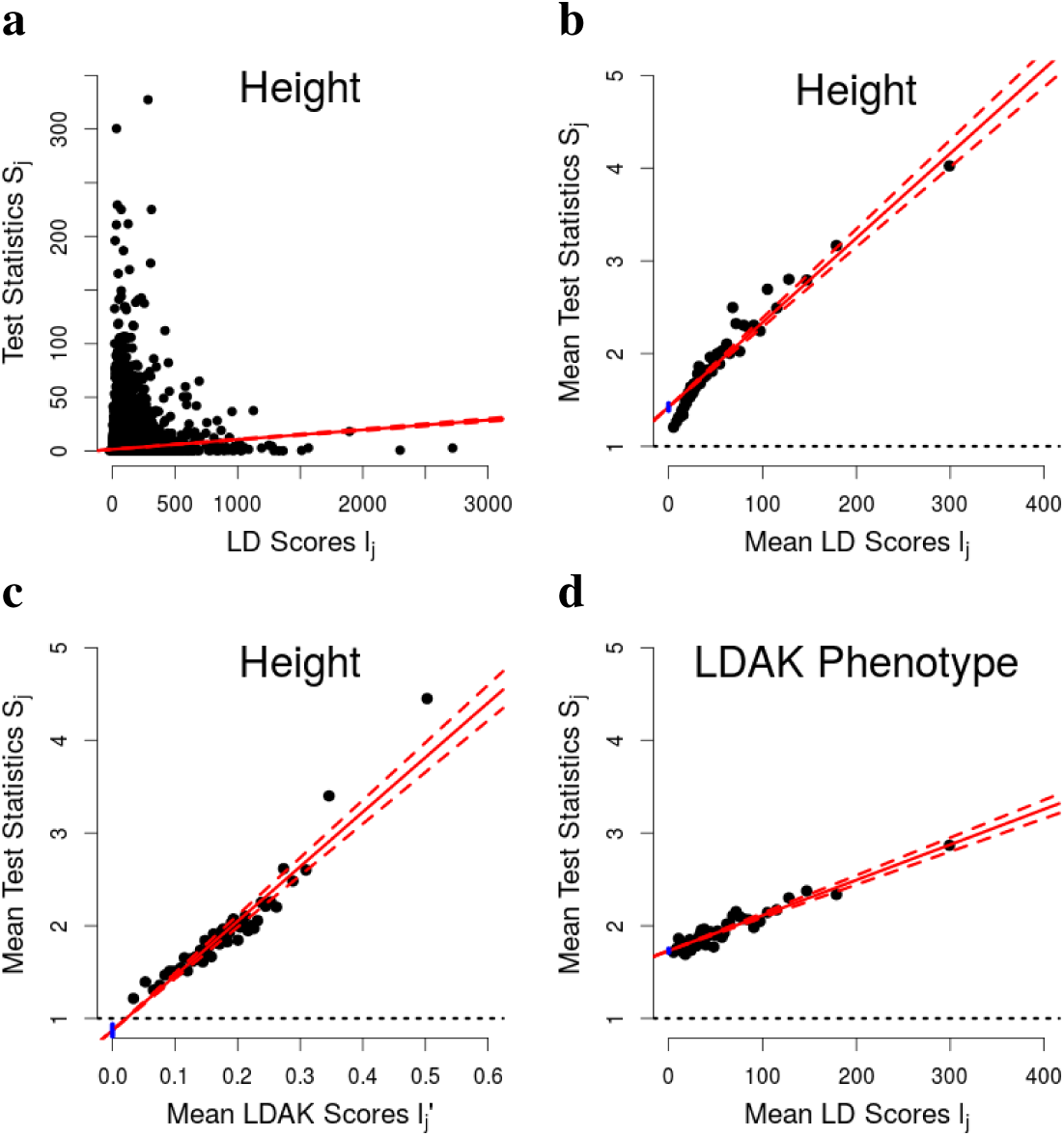
Test statistics are correlated with both LD and LDAK Scores. **(a)** Test statistics versus LD scores from the most recent Giant Consortium meta-analysis for height;^12^ to avoid correlated datapoints, we restrict to a subset of 121 310 SNPs with MAF*>*0.01 in approximate linkage equilibrium (obtained by pruning so that no two SNPs within 1 cM have 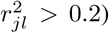). **(b)** The correlation can be magnified by first dividing SNPs into 50 bins based on LD Scores, then plotting mean test statistic versus mean LD score for each bin.^3^ **(c)** The same as (b), except we consider LDAK scores instead of LD Scores. **(d)** The same as (b), except that instead of using the test statistics for height, we generate new ones based on the LDAK Model. In each plot, the solid red line is the line of best fit from least-squares regression; the dashed red lines and solid blue segments indicate, respectively, 95% confidence intervals for the slope and intercept from this regression.

Secondly, a significant correlation between *l*_*j*_ and 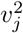 only proves that the GCTA Model fits better than the model 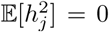, not that it fits better than the LDAK Model. The LDAK Model predicts 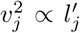; while Figure 1c shows that for height there is also significant correlation between 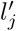 and *S*_*j*_, it would be equally absurd of us to claim that the LDAK Model was superior to the GCTA Model based on this evidence alone. For Figure 1d, we generate test statistics under the LDAK Model (assuming no confounding); specifically, we sample *S*_*j*_ from a 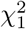 distribution with non-centrality parameter 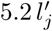 (we chose 5.2 so that the mean test statistic is 2.29, matching that observed for height). This simulation demonstrates that *l*_*j*_ and *S*_*j*_ will also be correlated for LDAK phenotypes, on account of the strong correlation between *l*_*j*_ and 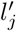 (for these data, their correlation is 0.51). Moreover, it highlights the dangers of using (S-)LDSC when the GCTA Model is not appropriate. The model used by LDSC makes strong predictions about how 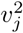, and therefore *S*_*j*_, vary across the genome; for example, the 95% range of *l*_*j*_ is 38 to 228, and the 10% (1%) of SNPs with highest *l*_*j*_ are on average expected to tag 2.8 (5.8) times as much heritability as the average SNP. When the data do not align with these predictions, LDSC will compensate by under-estimating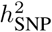 (the slope of the line) and over-estimating *a* (the intercept).

## Demonstrating problems with S-LDSC

The original S-LDSC model contained 53 categories: 28 functional annotations (which include coding, conserved and DHS regions), 24 buffers and the base category containing all SNPs.^6^ Recently, this was expanded to 75 categories, by adding 3 more functional annotations, 3 extra buffers, 10 MAF tranches and 6 continuous LD-related annotations.^7^ We now construct an additional category of “thinned SNPs”, by pruning so that no two SNPs within 1 cM have 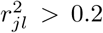, and also the corresponding buffer (all thinned SNPs and those within 500 bp). Table 1 and Supplementary Table 1 report average estimates of enrichment for coding, conserved, DHS and thinned SNPs, estimated using six versions of LDSC (which vary according to choice of category), as well as GCTA and LDAK. We use two sources of data: LDSC requires only summary statistics, so we first analyze published results from 24 large-scale GWAS (12 binary traits, 12 quantitative, average sample size 121 000; see Supplementary Table 2); GCTA and LDAK need raw data, so we also perform 25 GWAS using data from the Wellcome Trust Case Control Consortium^13^ and the eMerge Network^14^ (18 binary traits, 7 quantitative, average sample size 9 700; see Supplementary Table 3).

**Table 1:** Enrichment of coding, conserved, DHS and thinned SNPs. For each of the four annotations, values report average estimates of enrichment based on either summary statistics from 24 published GWAS, or analysis of 25 GWAS for which we have raw genotype and phenotype data. We use six versions of LDSC: 2-part (the annotation SNPs and the base category containing all SNPs); 3-part (the annotation SNPs, the corresponding 500 bp buffer and the base category); old S-LDSC (53 categories, including coding, conserved and DHS SNPs); old S-LDSC+ (the 53 categories, plus thinned SNPs and the corresponding buffer); new S-LDSC (75 categories); new S-LDSC+ (75 categories, plus thinned SNPs and the corresponding buffer). We also estimate enrichments using GCTA and LDAK, each time constructing two genomic similarity matrices (GSMs), the first corresponding to the annotation SNPs, the second to all other SNPs.

|  |  |  |  |  |  |
| --- | --- | --- | --- | --- | --- |
| Summary Statistics | 2-Part LDSC | 10.4 (0.5) | 18.1 (0.5) | 5.3 (0.1) | 24.2 (0.4) |
|  | 3-Part LDSC | 7.5 (0.5) | 10.6 (0.6) | 3.2 (0.1) | 24.6 (0.4) |
|  | Old S-LDSC | 6.2 (0.5) | 12.0 (0.5) | 1.7 (0.2) |  |
|  | Old S-LDSC+ | 4.6 (0.3) | 7.9 (0.3) | 1.5 (0.1) | 20.3 (0.4) |
|  | New S-LDSC | 4.5 (0.4) | 7.6 (0.4) | 1.4 (0.1) |  |
|  | New S-LDSC+ | 4.0 (0.3) | 6.3 (0.3) | 1.4 (0.1) | 14.5 (0.5) |
| Raw Data | 2-Part LDSC | 18.3 (1.7) | 18.3 (1.4) | 8.2 (0.2) | 28.7 (0.8) |
|  | 2-GRM GCTA | 15.3 (1.5) | 15.8 (1.3) | 7.6 (0.2) | 22.3 (0.9) |
|  | 2-GRM LDAK | 2.9 (0.4) | 1.9 (0.3) | 1.3 (0.1) | 0.9 (0.1) |

Table 1 highlights two shortcomings with using S-LDSC to estimate enrichments. Firstly, there are many arbitrary choices underlying S-LDSC, such as which functional categories and LD annotations to include, the size and number of buffer regions and how to partition SNPs by MAF; we see that estimates from S-LDSC vary substantially depending on these choices. Secondly, both old and new S-LDSC find thinned SNPs to be highly enriched for heritability: 20.3-fold (SD 0.4) and 14.5-fold (SD 0.5), respectively. Considering that we selected thinned SNPs simply by pruning, and not based on biological criteria, we see no reason why they should be many-fold enriched for heritability. By contrast, LDAK estimates their enrichment to be 0.86-fold (SD 0.06), indicating that the high estimates from S-LDSC are a consequence of the GCTA Model not accounting for LD.

In summary, Gazal *et al.* have argued that the heritability model used by LDSC better reflects real data than the LDAK Model, and that high estimates of functional enrichment from S-LDSC should be preferred to those from LDAK. We do not agree with their first claim; whereas we provided rigorous evidence to support the LDAK Model,^1^ Gazal *et al.* rely on the observation that for many traits, association test statistics correlate with LD scores, something we have shown is also to be expected under the LDAK Model. Nor do we agree with their second claim; we have shown that S-LDSC estimates can be highly sensitive to category choice, without it being clear which choice to prefer, and that simply by thinning SNPs, we can construct a category which S-LDSC finds to be over ten-fold enriched for heritability.

## URLs

LDAK, http://ldak.org; GCTA, http://cnsgenomics.com/software/gcta; LDSC, http://github.com/bulik/ldsc.

## Methods

Full details for repeating our analyses are provided in the Supplementary Note.

## Acknowledgments

Access to Wellcome Trust Case Control Consortium data was authorized as work related to the project “Genome-wide association study of susceptibility and clinical phenotypes in epilepsy,” access to eMerge Network data was granted under dbGaP Project 14422, “Comprehensive testing of SNP-based prediction models.” D.S. is funded by the UK Medical Research Council under grant MR/L012561/1, by the European Unions Horizon 2020 Research and Innovation Programme under the Marie Sklodowska-Curie grant agreement number 754513, and by Aarhus University Research Foundation (AUFF). The eMERGE Network was initiated and funded by NHGRI through the following grants: U01HG006828 (Cincinnati Childrens Hospital Medical Center/Boston Childrens Hospital); U01HG006830 (Childrens Hospital of Philadelphia); U01HG006389 (Essentia Institute of Rural Health, Marshfield Clinic Research Foundation and Pennsylvania State University); U01HG006382 (Geisinger Clinic); U01HG006375 (Group Health Cooperative); U01HG006379 (Mayo Clinic); U01HG006380 (Icahn School of Medicine at Mount Sinai); U01HG006388 (Northwestern University); U01HG006378 (Vanderbilt University Medical Center); and U01HG006385 (Vanderbilt University Medical Center serving as the Coordinating Center).

## Author contributions

D.S. performed the analysis, D.S. and D.J.B. wrote the manuscript.

## Competing financial interests

The authors declare no competing financial interests.

### Code availability

Step-by-step code for performing the analyses underlying Figure 1 and Table 1 are provided in the Supplementary Note.

### Data availability

WTCCC^13^ and eMerge Network^14^ were applied for and downloaded from the European Genome-phenome Archive and dbGaP, respectively (see Supplementary Table 3 for accession codes). Results for each of the 24 summary GWAS are available to download from the websites of the corresponding study (see Supplementary Table 2 for references).

## References

1. Speed, D. et al. Reevaluation of SNP heritability in complex human traits. Nat. Genet. 49, 986–992 (2017).

2. Yang, J., Lee, S., Goddard, M. & Visscher, P. GCTA: a tool for genome-wide complex trait analysis. Am. J. Hum. Genet. 88, 76–82 (2011).

3. Bulik-Sullivan, B. et al. LD score regression distinguishes confounding from polygenicity in genome-wide association studies. Nat. Genet. 47, 291–295 (2014).

4. Speed, D., Hemani, G., Johnson, M. & Balding, D. Improved heritability estimation from genome-wide SNP data. Am. J. Hum. Genet. 91, 1011–1021 (2012).

5. Gusev, A. et al. Partitioning Heritability of Regulatory and Cell-Type-Specific Variants across 11 Common Diseases. Am. J. Hum. Genet. 95, 535–552 (2014).

6. Finucane, H. et al. Partitioning heritability by functional annotation using genome-wide association summary statistics. Nat. Genet. 47, 1228–1235 (2015).

7. Gazal, S. et al. Linkage disequilibrium-dependent architecture of human complex traits shows action of negative selection. Nat. Genet. 49 (2017).

8. Yang, J. et al. Genomic inflation factors under polygenic inheritance. Eur. J. Hum. Genet. 19, 807–812 (2011).

9. Yang, J. et al. Genomic partitioning of genetic variation for complex traits using common SNPs. Nat. Genet. 43, 519–525 (2011).

10. Speed, D. et al. Describing the genetic architecture of epilepsy through heritability analysis. Brain 137, 26802689 (2014).

11. Yang, J. et al. Genetic variance estimation with imputed variants finds negligible missing heritability for human height and body mass index. Nat. Genet. 47, 1114–1120 (2015).

12. Wood, A. et al. Defining the role of common variation in the genomic and biological architecture of adult human height. Nat. Genet. 46, 1173–1186 (2014).

13. The Wellcome Trust Case Control Consortium. Genome-wide association study of 14,000 cases of seven common diseases and 3,000 shared controls. Nature 447, 661–678 (2007).

14. Verma, S. et al. Imputation and quality control steps for combining multiple genome-wide datasets. Front. Genet. 5, 370 (2015).

