## Supplementary Materials for "Exposing flaws in S-LDSC; reply to Gazal *et al*."

### Exposing flaws in S-LDSC; reply to Gazal *et al.* Supplementary Material

Doug Speed[1,2,*] and David J Balding.[2,3]

[1] *Aarhus Institute of Advanced Studies (AIAS), Aarhus University, Denmark.*
[2] *UCL Genetics Institute, University College London, United Kingdom.*
[3] *Melbourne Integrative Genomics, School of BioSciences and School of Mathematics & Statistics, University of Melbourne, Australia.*

**Supplementary Note: Details of analyses.** Here we provide step-by-step scripts for performing our two analyses, the one underlying Figure 1, which demonstrates that test statistics are correlated with both LD and LDAK scores, and the one underlying Table 1, which estimates enrichments using LDSC, GCTA and LDAK. These analyses require the softwares LDAK (available at http://ldak.org) and LDSC (available at http://github.com/bulik/ldsc); we assume that the LDAK executable is saved in current folder, so that LDAK can be run by typing `./ldak5.linux` (or `./ldak5.mac` for the Mac version), and that the LDSC files are stored in the folder `ldsc`, so that LDSC can be run by typing `python2 ldsc/ldsc.py`. Note that a backslash (\) at the end of a line indicates the command continues on the next line. This code is also available at http://ldak.org/protocol.

```
#PREPARATION

#Download and extract European 1000 Genomes Project Phase 3 data and frequencies
wget https://data.broadinstitute.org/alkesgroup/LDSCORE/1000G_Phase3_plinkfiles.tgz
wget https://data.broadinstitute.org/alkesgroup/LDSCORE/1000G_Phase3_frq.tgz
tar -xvzf 1000G_Phase3_plinkfiles.tgz
tar -xvzf 1000G_Phase3_frq.tgz

#The data are divided by chromosome; for convenience, rename these
for j in {1..22}; do
for end in {bed,bim,fam}; do mv 1000G_EUR_Phase3_plink/1000G.EUR.QC.$j.$end 1000G.$j.$end; done
mv 1000G_Phase3_frq/1000G.EUR.QC.$j.frq 1000G.$j.frq
done

#Download and extract LD scores for old S-LDSC (53 categories) and new S-LDSC (75 categories)
wget https://data.broadinstitute.org/alkesgroup/LDSCORE/1000G_Phase3_baseline_v1.1_ldscores.tgz
wget https://data.broadinstitute.org/alkesgroup/LDSCORE/1000G_Phase3_baselineLD_v1.1_ldscores.tgz
tar -xvzf 1000G_Phase3_baseline_v1.1_ldscores.tgz
tar -xvzf 1000G_Phase3_baselineLD_v1.1_ldscores.tgz

#Download HapMap3 SNPs
wget https://data.broadinstitute.org/alkesgroup/LDSCORE/w_hm3.snplist.bz2
bunzip2 w_hm3.snplist.bz2

#Download and tidy summary statistics; require name, alleles, Z stat, p-value and sample size (restrict to biallelic SNPs)
#Store summary statistics for trait # in #.prep

#This demonstrates for height (using results from GIANT Consortium)
wget https://portals.broadinstitute.org/collaboration/giant/images/0/01/\
GIANT_HEIGHT_Wood_et_al_2014_publicrelease_HapMapCeuFreq.txt.gz
gunzip -c GIANT_HEIGHT_Wood_et_al_2014_publicrelease_HapMapCeuFreq.txt.gz | \
awk '(NR==1){print "SNP","A1","A2","Z","P","N"}(NR>1 && ($2=="A"||$2=="C"||$2=="G"||$2=="T") \
&& ($3=="A"||$3=="C"||$3=="G"||$3=="T")){print $1, $2, $3, $5/$6, $7, $8}' - > height.prep

#FIGURE 1

#LD Scores for Chromosome # are provided in Column 4 of baseline_v1.1/baseline.#.l2.ldscore.gz
#Compute LDAK scores as follows
for j in {1..22}; do
./ldak5.linux --cut-weights chr$j --bfile 1000G.$j
./ldak5.linux --calc-weights-all chr$j --bfile 1000G.$j
./ldak5.linux --calc-tagging chr$j --bfile 1000G.$j --weights chr$j/weights.short \
--power -0.25 --window-cm 1 --reduce NO
done
#LDAK scores for Chromosome # are in Column 10 of chr#.tagging
```

```
#For plotting, require a pruned set of HapMap3 SNPs for which we have height test statistics
awk '(NR==FNR){arr[$1];next}($1 in arr){print $1, $4^2}' w_hm3.snplist height.prep > height.hapmap
mkdir plot
for j in {1..22}; do
./ldak5.linux --thin plot/chr$j --bfile 1000G.$j --window-prune .2 --window-cm 1 --extract height.hapmap
done
#Now collect up LD scores, LDAK scores and test statistics for pruned SNPs
rm plot.ld plot.ldak plot.stats
for j in {1..22}; do
gunzip -c baseline_v1.1/baseline.$j.l2.ldscore.gz | awk '(NR==FNR){arr[$1];next}($2 in arr){print $2, $4}' \
plot/chr$j.in - >> plot.ld
awk '(NR==FNR){arr[$1];next}($1 in arr){print $1, $10}' plot/chr$j.in chr$j.tagging >> plot.ldak
awk '(NR==FNR){arr[$1]=$2;next}{print $1, arr[$1]}' height.hapmap plot/chr$j.in >> plot.stats
done

#Can plot these in R
#To generate test statistics under the LDAK Model, use rchisq
#E.g, read the file plot.ldak into the matrix ldak, then use rchisq(nrow(ldak),1,ncp=5.2*ldak[,2])

#TABLE 1

#First, we analyzed summary statistics from 24 public GWAS using six versions of LDSC

#For each trait, remove MHC, SNPs with test statistic > max(80,n/10000), then "munge" to get in LDSC format
#If info scores are provided, also exclude SNPs with score <.95
#Summary statistics for trait # will be stored in #.sumstats.gz

#This demonstrates for height (n=253288)
awk < 1000G.6.bim '($4>=25000000 && $4<=34000000){print $2}' > mhc.snps
awk -v n=253288 '(NR==FNR){arr[$1];next}($1 in arr || ($4^2>n/10000 && $4^2>80)){print $1}' mhc.snps height.prep > excl.snps
awk '(NR==FNR){arr[$1];next}!($1 in arr){print $0}' excl.snps height.prep > height.stats
python2 ldsc/munge_sumstats.py --out height --sumstats height.stats --merge-alleles w_hm3.snplist

#Need LD scores for the base category (all SNPs) - can extract these from the 53-category LD scores
mkdir base
for j in {1..22}; do
gunzip -c baseline_v1.1/baseline.$j.annot.gz | awk '{print $1, $2, $3, $4, $5}' - | gzip > base/chr$j.annot.gz
gunzip -c baseline_v1.1/baseline.$j.l2.ldscore.gz | awk '{print $1, $2, $3, $4}' - | gzip > base/chr$j.l2.ldscore.gz
awk '{print $1}' baseline_v1.1/baseline.$j.l2.M > base/chr$j.l2.M
awk '{print $1}' baseline_v1.1/baseline.$j.l2.M_5_50 > base/chr$j.l2.M_5_50
done

#and likewise for coding SNPs (annotation 2 of 53)
mkdir coding
k=2
for j in {1..22}; do
gunzip -c baseline_v1.1/baseline.$j.annot.gz | awk -v k=$k '{print $1,$2,$3,$4,$(4+k)}' - | gzip > coding/chr$j.annot.gz
gunzip -c baseline_v1.1/baseline.$j.l2.ldscore.gz | awk -v k=$k '{print $1,$2,$3,$(3+k)}' - | gzip > coding/chr$j.l2.ldscore.gz
awk -v k=$k '{print $k}' baseline_v1.1/baseline.$j.l2.M > coding/chr$j.l2.M
awk -v k=$k '{print $k}' baseline_v1.1/baseline.$j.l2.M_5_50 > coding/chr$j.l2.M_5_50
done

#and for coding SNPs buffer (annotation 3 of 53)
mkdir codingb
k=2
for j in {1..22}; do
gunzip -c baseline_v1.1/baseline.$j.annot.gz | awk -v k=$k '{print $1,$2,$3,$4,$(4+k)}' - | gzip > codingb/chr$j.annot.gz
gunzip -c baseline_v1.1/baseline.$j.l2.ldscore.gz | awk -v k=$k '{print $1,$2,$3,$(3+k)}' - | gzip > codingb/chr$j.l2.ldscore.gz
awk -v k=$k '{print $k}' baseline_v1.1/baseline.$j.l2.M > codingb/chr$j.l2.M
awk -v k=$k '{print $k}' baseline_v1.1/baseline.$j.l2.M_5_50 > codingb/chr$j.l2.M_5_50
done

#Repeat this for conserved SNPs, their buffer, DHS SNPs and their buffer (annotations 4, 5, 11 and 12)
#Save these in folders conserved, conservedb, dhs and dhsb

#Make annotation files for thinned SNPs and a 500bp buffer
mkdir thinned thinnedb
for j in {1..22}; do
./ldak5.linux --thin thinned/chr$j --bfile 1000G.$j --window-prune .2 --window-cm 1
gunzip -c baseline_v1.1/baseline.$j.annot.gz | awk '(NR==FNR){arr[$1];next}(FNR==1){print "CHR","BP","SNP","CM","THINNED"}\
(FNR>1){thinned=0;if($3 in arr){thinned=1}{print $1, $2, $3, $4, thinned}' thinned/chr$j.in - | gzip > thinned/chr$j.annot.gz
```

```
awk '(NR==FNR){arr[$1];next}($2 in arr){a++;print a, $1, $4, $4}' thinned/chr$j.in 1000G.$j.bim > thinnedb/chr$j.regions
ldak5.linux --cut-genes temp$j --bfile 1000G.$j --genefile thinnedb/chr$j.regions --gene-buffer 500 --ignore-weights YES
mv temp$j/genes.predictors.used thinnedb/chr$j.in
gunzip -c baseline_v1.1/baseline.$j.annot.gz | awk '(NR==FNR){arr[$1];next}(FNR==1){print "CHR","BP","SNP","CM","BUFFER"}\
(FNR>1){thinned=0;if($3 in arr){thinned=1}{print $1, $2, $3, $4, thinned}' thinnedb/chr$j.in - | gzip > thinnedb/chr$j.annot.gz
done
#and compute LD scores for these two annotations
for j in {1..22}; do
python2 ldsc/ldsc.py --bfile 1000G.$j --ld-wind-cm 1 --l2 --out thinned/chr$j
--annot thinned/chr$j.annot.gz --print-snps baseline_v1.1/baseline.$j.snps
python2 ldsc/ldsc.py --bfile 1000G.$j --ld-wind-cm 1 --l2 --out thinnedb/chr$j \
--annot thinnedb/chr$j.annot.gz --print-snps baseline_v1.1/baseline.$j.snps
done

#Now estimate enrichments for each trait
trait=height  #this determines which trait to test

#Get LD scores for the SNPs with summary statistics (the regression weights)
mkdir weights$trait
gunzip -c $trait.sumstats.gz | awk '(NR>1){print $1}' > weights$trait/snps
for j in {1..22}; do
python2 ldsc/ldsc.py --bfile 1000G.$j --ld-wind-cm 1 --l2 --out weights$trait/chr$j --extract weights$trait/snps
done

#Do 2- and 3-part LDSC for coding, conserved, dhs and thinned SNPs
for name in {coding,conserved,dhs,thinned}; do
python2 ldsc/ldsc.py --h2 $trait.sumstats.gz --w-ld-chr weights$trait/chr --overlap-annot --frqfile-chr 1000G. \
--ref-ld-chr base/chr,$name/chr --out res$trait.2.part.$name --print-coefficients
python2 ldsc/ldsc.py --h2 $trait.sumstats.gz --w-ld-chr weights$trait/chr --overlap-annot --frqfile-chr 1000G. \
--ref-ld-chr base/chr,$name/chr,${name}b/chr --out res$trait.3.part.$name --print-coefficients
done

#Old S-LDSC and old S-LDSC+
python2 ldsc/ldsc.py --h2 $trait.sumstats.gz --w-ld-chr weights$trait/chr --overlap-annot --frqfile-chr 1000G. \
--ref-ld-chr baseline_v1.1/baseline. --out res$trait.53.part --print-coefficients
python2 ldsc/ldsc.py --h2 $trait.sumstats.gz --w-ld-chr weights$trait/chr --overlap-annot --frqfile-chr 1000G. \
--ref-ld-chr baseline_v1.1/baseline.,thinned/chr,thinnedb/chr --out res$trait.55.part --print-coefficients

#New S-LDSC and new S-LDSC+
python2 ldsc/ldsc.py --h2 $trait.sumstats.gz --w-ld-chr weights$trait/chr --overlap-annot --frqfile-chr 1000G. \
--ref-ld-chr baselineLD_v1.1/baselineLD. --out res$trait.75.part --print-coefficients
python2 ldsc/ldsc.py --h2 $trait.sumstats.gz --w-ld-chr weights$trait/chr --overlap-annot --frqfile-chr 1000G. \
--ref-ld-chr baselineLD_v1.1/baselineLD.,thinned/chr,thinnedb/chr --out res$trait.77.part --print-coefficients

#After testing each trait, can combine (meta-analyze) estimates across traits using commands such as
grep Coding_UCSC.bedL2_0 res*.53.part.results | awk '{a+=$5/$6^2;b+=1/$6^2}END{print NR, "Estimate:",a/b, "SD:",1/b^.5}'

#Second, we applied LDSC, GCTA LDAK to raw data for 25 traits

#Suppose (post-QC) genotypes are stored in data.bed, data.bim and data.fam, and phenotypes in phen.pheno
#The file excl.snps contains SNPs to exclude (those in MHC or with test statistic >80)
#Also have a covariate file, cov.covar, containing sex, 20 PCs from data, and 10 projections from 1000 Genomes
#For details of QC and for making this covariate file, see http://dougspeed.com/protocol ("Merging Cohorts")

#Prune to obtain a list of thinned SNPs
./ldak5.linux --thin thinned --bfile data --window-prune .2 --window-cm 1

#Find which SNPs are within coding, conserved and DHS regions
wget https://data.broadinstitute.org/alkesgroup/LDSCORE/baseline_bedfiles.tgz
tar -xvzf baseline_bedfiles.tgz

rm coding.regions
for j in {1..22}; do awk -v j=$j '($1=="chr"j){print NR,j,$2,$3}' baseline/Coding_UCSC.bed >> coding.regions; done
rm conserved.regions
for j in {1..22}; do awk -v j=$j '($1=="chr"j){print NR,j,$2,$3}' baseline/Conserved_LindbladToh.bed >> conserved.regions; done
rm dhs.regions
for j in {1..22}; do awk -v j=$j '($1=="chr"j){print NR,j,$2,$3}' baseline/DHS_peaks_Trynka.bed >> dhs.regions; done

./ldak5.linux --cut-genes tempcoding --bfile data --genefile coding.regions --ignore-weights YES
mv tempcoding/genes.predictors.used coding.in
./ldak5.linux --cut-genes tempconserved --bfile data --genefile conserved.regions --ignore-weights YES
```

```
mv tempconserved/genes.predictors.used conserved.in
./ldak5.linux --cut-genes tempdhs --bfile data --genefile dhs.regions --ignore-weights YES
mv tempdhs/genes.predictors.used dhs.in

#The steps for 2-part LDSC are as above, except we used data.bed, data.bim and data.fam as the reference panel (not 1000 Genomes)

#For GCTA and LDAK, we start by constructing genome-wide kinship matrices
./ldak5.linux --cut-weights sections --bfile data
./ldak5.linux --calc-weights-all sections --bfile data
./ldak5.linux --calc-kins-direct ldak_all --bfile data --weights sections/weights.short --power -.25 --exclude excl.snps
./ldak5.linux --calc-kins-direct gcta_all --bfile data --ignore-weights YES --power -1 --exclude excl.snps

#Now construct kinship matrices for each annotation, and their complements
for name in {coding,conserved,dhs,thinned}; do
./ldak5.linux --calc-kins-direct ldak_$name --bfile data --weights sections/weights.short --power -.25 \
--exclude excl.snps --extract $name.in
echo "ldak_all
ldak_$name" > sub.txt
./ldak5.linux --sub-grm ldak_not_$name --mgrm sub.txt
./ldak5.linux --calc-kins-direct gcta_$name --bfile data --ignore-weights YES --power -1 \
--exclude excl.snps --extract $name.in
echo "gcta_all
gcta_$name" > sub.txt
./ldak5.linux --sub-grm gcta_not_$name --mgrm sub.txt
done

#Finish by performing REML for each annotation
for name in {coding,conserved,dhs,thinned}; do
echo "gcta_$name
gcta_not_$name" > mlist.txt
./ldak5.linux --reml reml_gcta_$name --covar cov.covar --pheno phen.pheno --mgrm mlist.txt
echo "ldak_$name
ldak_not_$name" > mlist.txt
./ldak5.linux --reml reml_ldak_$name --covar cov.covar --pheno phen.pheno --mgrm mlist.txt
done
#The enrichments are stored in the files reml_gcta_$name.share and reml_ldak_$name.share
```

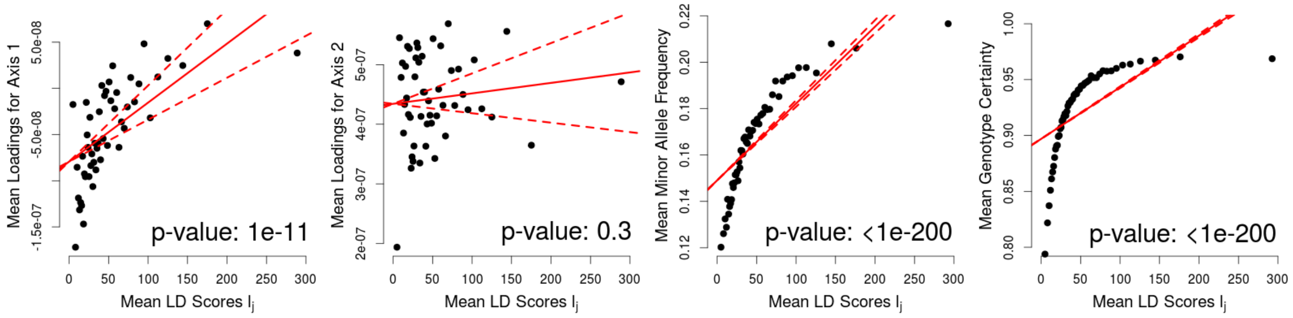


**Supplementary Figure 1: LD scores are correlated with population axes, minor allele frequencies and genotype certainties.** LDSC assumes that $a_j$, the inflation of test statistics due to confounding, is constant across SNPs,[1] and therefore orthogonal to LD scores, $l_j$. While a theoretical justification was provided for this assumption, it was not tested empirically. These plots demonstrate that $l_j$ is significantly correlated with population loadings, minor allele frequencies (MAF) and genotype certainties. Therefore it seems plausible that $l_j$ also correlates with $a_j$, violating the assumption of LDSC. For this figure, we restrict to 126 214 HapMap 3 SNPs[2] with MAF >0.01 and in approximate linkage equilibrium (obtained by pruning so that no two SNPs within 1 cM have $r^2_{jl} > 0.2$). For each plot, we divide the SNPs into 50 bins based on LD scores, then report means for each bin. LD scores are estimated using 489 European individuals and 9 997 231 SNPs from the 1000 Genomes Project.[3] The population loadings are obtained by performing principal component analysis using the 1 184 HapMap 3 individuals; here we consider the first two axes, which predominantly capture global population structure. The MAFs are estimated from the 489 European 1000 Genomes individuals. As a proxy for genotype certainties, we use the info scores reported by the most recent Psychiatric Genomics Consortium meta-analysis for schizophrenia.[4] In each plot, the solid red line is the line of best fit from least-squares regression, while the dashed red lines indicate 95% confidence intervals for the slope from this regression (the $p$-value corresponds to testing whether the slope is 0).

| | | Average Estimated Percentage of $h^2_{\text{SNP}}$ (SD) | | | | Expected Percentage of $h^2_{\text{SNP}}$ (SD) | | | | Average Estimated Enrichment (SD) | | | |
|---|---|---|---|---|---|---|---|---|---|---|---|---|---|
| | | **Coding** | **Conserved** | **DHS** | **Thinned** | **Coding** | **Conserved** | **DHS** | **Thinned** | **Coding** | **Conserved** | **DHS** | **Thinned** |
| Summary Statistics | 2-Part LDSC | 14.8 (0.7) | 46.5 (1.2) | 88.8 (1.4) | 51.3 (0.8) | 1.4 | 2.6 | 16.6 | 2.1 | 10.4 (0.5) | 18.1 (0.5) | 5.3 (0.1) | 24.2 (0.4) |
| | 3-Part LDSC | 10.7 (0.8) | 27.3 (1.4) | 53.4 (2.5) | 52.4 (0.8) | 1.4 | 2.6 | 16.6 | 2.1 | 7.5 (0.5) | 10.6 (0.6) | 3.2 (0.1) | 24.6 (0.4) |
| | Old S-LDSC | 8.8 (0.7) | 30.7 (1.2) | 28.1 (2.6) | | 1.4 | 2.6 | 16.6 | | 6.2 (0.5) | 12.0 (0.5) | 1.7 (0.2) | |
| | Old S-LDSC+ | 6.5 (0.4) | 20.4 (0.8) | 25.2 (1.7) | 43.2 (0.8) | 1.4 | 2.6 | 16.6 | 2.1 | 4.6 (0.3) | 7.9 (0.3) | 1.5 (0.1) | 20.3 (0.4) |
| | New S-LDSC | 6.5 (0.6) | 19.5 (1.1) | 23.1 (2.3) | | 1.4 | 2.6 | 16.6 | | 4.5 (0.4) | 7.6 (0.4) | 1.4 (0.1) | |
| | New S-LDSC+ | 5.7 (0.5) | 16.2 (0.9) | 22.7 (1.9) | 30.8 (1.1) | 1.4 | 2.6 | 16.6 | 2.1 | 4.0 (0.3) | 6.3 (0.3) | 1.4 (0.1) | 14.5 (0.5) |
| Raw Data | 2-Part LDSC | 22.4 (2.0) | 46.3 (3.5) | 126.1 (2.5) | 74.5 (2.1) | 1.2 | 2.5 | 15.4 | 2.6 | 18.3 (1.7) | 18.3 (1.4) | 8.2 (0.2) | 28.7 (0.8) |
| | 2-GRM GCTA | 18.7 (1.8) | 40.0 (3.3) | 117.4 (3.3) | 58.2 (2.4) | 1.2 | 2.5 | 15.4 | 2.6 | 15.3 (1.5) | 15.8 (1.3) | 7.6 (0.2) | 22.3 (0.9) |
| | 2-GRM LDAK | 4.3 (0.6) | 5.7 (0.8) | 23.2 (1.8) | 14.1 (1.5) | 1.5 | 2.9 | 17.8 | 16.5 | 2.9 (0.4) | 1.9 (0.3) | 1.3 (0.1) | 0.9 (0.1) |

**Supplementary Table 1: Enrichment of coding, conserved, DHS and thinned SNPs.** This is an enlarged version of Table 1 in the main text. Values report the average percentage of $h^2_{\text{SNP}}$ that each annotation is estimated to contribute, the percentage of $h^2_{\text{SNP}}$ that each is expected to contribute, and the average estimated enrichment of each (the estimated percentage divided by the expected percentage). Estimates are obtained either using summary statistics from 24 published GWAS (Supplementary Table 2), or from analysis of 25 GWAS for which we have raw genotype and phenotype data (Supplementary Table 3). We use six versions of LDSC, which vary according to the choice of categories: 2-part (the annotation SNPs and the base category containing all SNPs); 3-part (the annotation SNPs, the corresponding 500 bp buffer and the base category); old S-LDSC (53 categories, including coding, conserved and DHS SNPs); old S-LDSC+ (the 53 categories, plus thinned SNPs and the corresponding buffer); new S-LDSC (75 categories); new S-LDSC+ (75 categories, plus thinned SNPs and the corresponding buffer). We also estimate enrichments using GCTA and LDAK, each time constructing two genomic similarity matrices (GSMs), the first corresponding to the annotation SNPs, the second to all other SNPs.

| **Binary Traits** | **Average Sample Size** | **Quantitative Traits** | **Average Sample Size** |
|---|---|---|---|
| Alzheimer's Diseases[5] | 54 000 | Bone Mineral Density[6] | 33 000 |
| Coronary Artery[7] | 80 000 | Body Mass Index[8] | 229 000 |
| Crohn's Disease[9] | 21 000 | Fasting Glucose[10] | 58 000 |
| Depression[11] | 161 000 | Glycated Hemoglobin[12] | 46 000 |
| Ever Smoked?[13] | 74 000 | HDL Cholesterol[14] | 95 000 |
| Inflammatory Bowel[9] | 35 000 | Height[15] | 245 000 |
| Neuroticism[11] | 171 000 | LDL Cholesterol[14] | 90 000 |
| Rheumatoid Arthritis[16] | 58 000 | Menarche Age[17] | 253 000 |
| Schizophrenia[4] | 81 000 | Menopause Age[18] | 69 000 |
| Subjective-Wellbeing[11] | 298 000 | Triglyceride Levels[14] | 92 000 |
| Type 2 Diabetes[19] | 157 000 | Waist-Hip Ratio[20] | 141 000 |
| Ulcerative Colitis[9] | 27 000 | Years Education[21] | 329 000 |
| **Average** | **101 000** | **Average** | **140 000** |

**Supplementary Table 2: Details of the 24 summary traits.** This table lists the 12 binary and 12 quantitative GWAS for which we obtained summary statistics, as well as the average sample size of each (or the total sample size, if per-SNP sample sizes were not provided). Our search for publicly-available summary statistics was based on Table 1 of Pasaniuc and Price,[22] which provides links to GWAS results for 25 traits. Of these, we used 23, excluding hip and waist circumference due to their overlap with with waist-hip ratio; when multiple GWAS were listed for a trait, we picked the one with largest sample size. The 24th trait was menopause age, which we found by searching LD Hub.[23] All GWAS were predominantly European. For each trait, we considered only biallelic SNPs, and if info scores were available, we excluded SNPs with score <0.95. As recommended by LDSC,[1] we restricted to HapMap 3 SNPs,[2] and omitted SNPs within the major histocompatibility complex (Chromosome 6: 25-34 Mb), or with test statistic greater than max(80,$n$/10000), where $n$ is the total sample size of the GWAS.

| WTCCC | Sample Size | Number of SNPs | eMerge Network | Sample Size | Number of SNPs |
|---|---|---|---|---|---|
| Bipolar Disorder | 4788 | 1937638 | Age-related Macular Disease | 8475 | 2972162 |
| Coronary Artery Disease | 4857 | 1943684 | Heart Failure | 9005 | 2972162 |
| Crohn's Disease | 4653 | 1932019 | Peripheral Arterial Disease | 10541 | 2972162 |
| Hypertension | 4865 | 1944541 | Shingles (Herpes Zoster) | 11966 | 2972162 |
| Rheumatoid Arthritis | 4781 | 1941327 | Venous Thromboembolism | 13961 | 2972162 |
| Type I Diabetes | 4890 | 1937840 | Triglyceride | 12137 | 2972162 |
| Type II Diabetes | 4841 | 1940920 | LDL Cholesterol | 13420 | 2972162 |
| Barrett's Oesophagus | 7049 | 2887629 | HDL Cholesterol | 13788 | 2972162 |
| Celiac Disease | 9946 | 2876170 | Systolic Blood Pressure | 15058 | 2972162 |
| Ischaemic Stroke | 8982 | 2879412 | Bystolic Blood Pressure | 15062 | 2972162 |
| Parkinson's Disease | 6871 | 2877852 | Height | 18152 | 2972162 |
| Psoriasis | 7474 | 1685001 | Body Mass Index | 19309 | 2972162 |
| Ulcerative Colitis | 8020 | 1875844 | | | |
| **Average** | **6309** | **2204606** | **Average** | **13406** | **2972162** |

**Supplementary Table 3: Details of the 25 Raw GWAS.** We performed 13 GWAS using data from the Wellcome Trust Case Control Consortium[24] (WTCCC) and 12 GWAS using data from the eMERGE Network.[25] A full description of the 13 WTCCC GWAS is provided in our previous publication,[26] which includes details of phenotyping, assession codes, imputation and our strict quality control steps. The eMerge data was obtained from dbGaP (assession codes phs000888.v1.p1.c1, phs000888.v1.p1.c3, phs000888.v1.p1.c4 and phs000888.v1.p1.c5). These data were provided post-imputation, but thereafter we performed the same quality control steps as for the WTCCC GWAS (in summary, we excluded individuals with extreme missingness or heterozygosity, and those inferred to be of non-European ethnicity, and excluded SNPs with MAF <0.01, call rate <0.95 and info score <0.99).

WTCCC data were obtained from https://ebi.ac.uk/ega; the accession codes are EGAD00000000001, EGAD00000000002, EGAD00000000003, EGAD00000000004, EGAD00000000005, EGAD00000000006, EGAD00000000007, EGAD00000000008, EGAD00000000009 (WTCCC 1 studies) and EGAD00000000021, EGAD00000000022, EGAD00000000023, EGAD00000000024, EGAD00000000025, EGAD00000000057, EGAD00010000124, EGAD00010000264, EGAD00010000506, EGAD00010000634, EGAS00001000672 (WTCCC 2 studies). eMERGE Network data were obtained from https://ncbi.nlm.nih.gov/gap; the accession codes are phs000888.v1.p1.c1, phs000888.v1.p1.c3, phs000888.v1.p1.c4, phs000888.v1.p1.c5.


1. Bulik-Sullivan, B. *et al.* LD score regression distinguishes confounding from polygenicity in genome-wide association studies. *Nat. Genet.* **47**, 291–295 (2014).

2. The International HapMap 3 Consortium. Integrating common and rare genetic variation in diverse human populations. *Nature* **467**, 52–58 (2010).

3. The 1000 Genomes Project Consortium. A map of human genome variation from population-scale sequencing. *Nature* **467**, 1061–1073 (2010).

4. Schizophrenia Working Group of the Psychiatric Genomics Consortium. Biological insights from 108 schizophrenia-associated genetic loci. *Nature* **511**, 421–427 (2014).

5. Lambert, J. *et al.* Meta-analysis of 74,046 individuals identifies 11 new susceptibility loci for Alzheimer's disease. *Nat. Genet.* **45**, 1452–1458 (2013).

6. Zheng, H. *et al.* Whole-genome sequencing identifies EN1 as a determinant of bone density and fracture. *Nature* **526**, 112–117 (2015).

7. Schunkert, H. *et al.* Large-scale association analysis identifies 13 new susceptibility loci for coronary artery disease. *Nat. Genet.* **43**, 333–338 (2011).

8. Locke, A. *et al.* Genetic studies of body mass index yield new insights for obesity biology. *Nature* **518**, 197–206 (2015).

9. Liu, J. *et al.* Association analyses identify 38 susceptibility loci for inflammatory bowel disease and highlight shared genetic risk across populations. *Nat. Genet.* **47**, 979–986 (2015).

10. Manning, A. *et al.* A genome-wide approach accounting for body mass index identifies genetic variants influencing fasting glycemic traits and insulin resistance. *Nat. Genet.* **44**, 659–669 (2012).

11. Okbay, A. *et al.* Genetic variants associated with subjective well-being, depressive symptoms, and neuroticism identified through genome-wide analyses. *Nat. Genet.* **48**, 626–633 (2016).

12. Soranzo, N. *et al.* Common variants at 10 genomic loci influence hemoglobin a(c) levels via glycemic and nonglycemic pathway. *Diabetes* **59**, 3229–3239 (2010).

13. The Tobacco and Genetics Consortium. Genome-wide meta-analyses identify multiple loci associated with smoking behavior. *Nat. Genet.* **42**, 441–447 (2010).

14. Global Lipids Genetics Consortium. Discovery and refinement of loci associated with lipid levels. *Nat. Genet.* **45**, 1274–1283 (2013).

15. Wood, A. *et al.* Defining the role of common variation in the genomic and biological architecture of adult human height. *Nat. Genet.* **46**, 1173–1186 (2014).

16. Okada, Y. *et al.* Genetics of rheumatoid arthritis contributes to biology and drug discovery. *Nature* **506**, 376–381 (2014).

17. Perry, J. *et al.* Parent-of-origin-specific allelic associations among 106 genomic loci for age at menarche. *Nature* **514**, 92–97 (2014).

18. Day, F. *et al.* Large-scale genomic analyses link reproductive aging to hypothalamic signaling, breast cancer susceptibility and brca1-mediated dna repair. *Nat. Genet.* **47**, 1294–1303 (2015).

19. Scott, R. *et al.* An expanded genome-wide association study of type 2 diabetes in Europeans. *Diabetes* **66**, 2888–2902 (2017).

20. Shungin, D. *et al.* New genetic loci link adipose and insulin biology to body fat distribution. *Nat. Genet.* **518**, 187–196 (2015).

21. Okbay, A. *et al.* Genome-wide association study identifies 74 loci associated with educational attainment. *Nature* **533**, 539–542 (2016).

22. Pasaniuc, B. & Price, A. Dissecting the genetics of complex traits using summary association statistics. *Nat. Rev. Genet.* **18**, 117–127 (2017).

23. Zheng, J. *et al.* LD Hub: a centralized database and web interface to perform ld score regression that maximizes the potential of summary level GWAS data for SNP heritability and genetic correlation analysis. *Bioinformatics* **33**, 272–279 (2016).

24. The Wellcome Trust Case Control Consortium. Genome-wide association study of 14,000 cases of seven common diseases and 3,000 shared controls. *Nature* **447**, 661–678 (2007).

25. Verma, S. *et al.* Imputation and quality control steps for combining multiple genome-wide datasets. *Front. Genet.* **5**, 370 (2015).

26. Speed, D. *et al.* Reevaluation of SNP heritability in complex human traits. *Nat. Genet.* **49**, 986–992 (2017).
